# ClockstaRX: testing molecular clock hypotheses with genomic data

**DOI:** 10.1101/2023.02.02.526226

**Authors:** David A. Duchêne, Sebastián Duchêne, Josefin Stiller, Rasmus Heller, Simon Y. W. Ho

## Abstract

Phylogenetic studies of genomic data can provide valuable opportunities for evaluating evolutionary timescales and drivers of rate variation. These analyses require statistical tools based on molecular clocks. We present ClockstaRX, a flexible platform for exploring and testing evolutionary rate signals in phylogenomic data. It implements methods that use information from gene trees in Euclidean space, allowing data transformation, visualization, and hypothesis testing. ClockstaRX implements formal tests of the dimensionality reducibility of the Euclidean space of rates, and for identifying loci and branches that have a large influence on rate variation. Using simulations to evaluate the performance of the methods implemented, we find that inferences about rates can be strongly influenced by the overall amount of rate variation in the data, the shared patterns of among-lineage rate heterogeneity across groups of loci, and missing data. In an analysis of phylogenomic data from birds, we find a higher rate of evolution in introns compared with exons across all lineages. In addition, passerine taxa are highlighted as having unique patterns of genomic evolutionary rates compared with other avian lineages. Drawing on these results, we recommend careful exploratory analyses and filtering before performing phylogenomic analyses using molecular clocks.

## Introduction

The molecular clock provides the foundation for studies of evolutionary rates and timescales. Molecular clock models that can account for rate variation across genes and lineages are now routinely implemented in phylogenetic analyses (Ho and Duchêne 2014; dos Reis et al. 2016). The biological causes of variation in evolutionary rates include factors associated with the environment (Gillman et al. 2009), life history (Bromham 2009; Iglesias-Carrasco et al. 2019), and functional groups of genes (Drummond and Wilke 2008; Yang and Gaut 2011). Phylogenomic data offer valuable opportunities to understand the drivers of this variation in evolutionary rates, but their complexity and large size call for dedicated tools for visualization and model comparison.

To study evolutionary rate variation, it is convenient to partition the biological influences on rates into lineage effects, gene effects, and residual effects (Gillespie 1991; Gaut et al. 2011). Lineage effects are those that drive rate change across the whole genome in a given lineage (Bromham 2011), such as generation time (e.g., Hua et al. 2015) or metabolic rate (e.g., Montoya et al. 2022). Gene effects are those that lead to variation in rates across loci, as in the case of differing selective constraints among coding and non-coding DNA (e.g., Hughes and Yeager 1997; Laroche et al. 1997). Lastly, residual effects are formed by the interaction between gene effects and lineage effects (Takahata 1987; Cutler 2000; Bedford and Hartl 2008). For instance, a subset of loci might experience a new selective constraint and consequent reduction in evolutionary rate in a subset of the lineages being studied.

Disentangling the various forms of evolutionary rate variation is a difficult exercise, particularly for genome-scale data. Challenges include describing high-dimensional phylogenetic data (D. A. Duchêne et al. 2018; Smith 2021), accounting for gene-tree discordance (Mendes and Hahn 2016), and formulating meaningful biological hypotheses (Smith and Eyre-Walker 2003). One approach to the statistical analysis of rate variation is through model comparison. A simple model is the ‘universal pacemaker’ (Wolf et al. 2009; Snir et al. 2012), whereby all loci follow the same pattern of among-lineage rate variation. In this model, locus rates are fully correlated and can be described as a single variable or dimension that represents their relative rates (Fig. 1a). The relative rates might reflect the differences in selection coefficient across loci, while among-lineage rate variation might be explained by general species traits, such as generation time. A slight extension of this model is a ‘local pacemaker’ model stating that some lineages have greater variance than others in their rates across loci, in addition to loci having different relative rates (Fig. 1b). These patterns would reflect differences among lineages in factors such as selection coefficients or DNA repair efficiency.

**Fig. 1.**
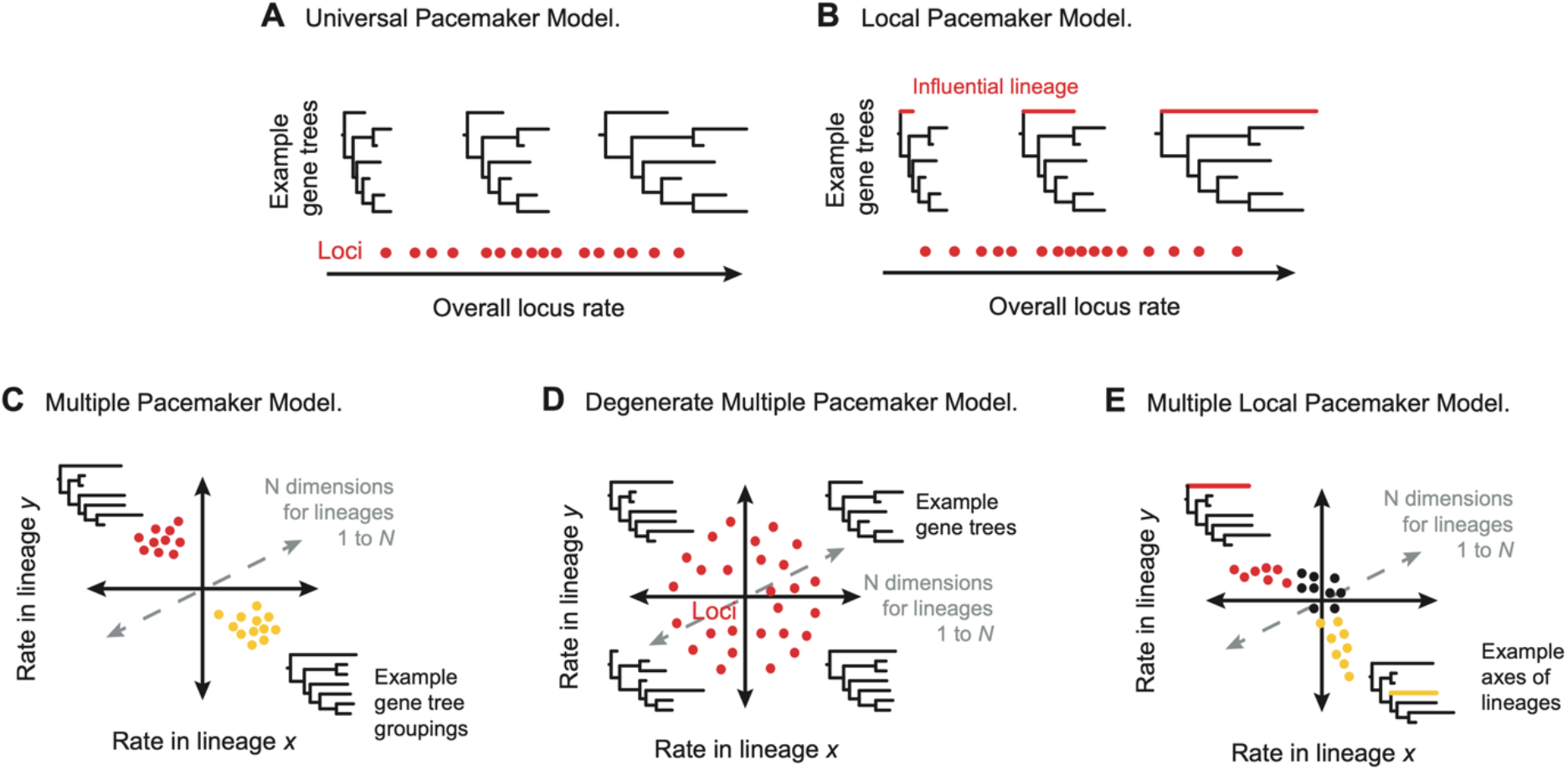
Hypotheses that can be evaluated in a Euclidean space of evolutionary rates with *N* dimensions for branches in a species tree. (a) Universal pacemaker model. Loci have evolutionary rates that co-vary across lineages, such that all variation can be explained by a single summary dimension that minimizes the variance across Euclidean space. (b) Local pacemaker model. A subset of lineages has a disproportionate influence on the overall variation in rates, such that they have high variance across the summarized space. (c) Multiple pacemaker model. Loci share patterns of among-lineage rate heterogeneity that are distinct from those of other groups of loci, such that these loci might group across space. (d) Degenerate multiple pacemaker model. Each locus has an independent pattern of among-lineage rate heterogeneity, such that the variables are completely uncorrelated. This model can serve as a null hypothesis. (e) Multiple local pacemaker model. Loci vary in their contribution to the variance in subsets of lineages, such that they follow a continuous trend across space.

The ‘multiple pacemaker’ model is of intermediate complexity and states that loci can be grouped into discrete bins of among-lineage rate variation (Snir et al. 2012; Duchêne and Ho 2015). This might be due to a set of loci experiencing a shared selective pressure that induces concomitant changes in their evolutionary rates (Fig. 1c). These groups of loci can be examined for enriched functional terms that might have been adaptively important for the taxa involved. The groupings can also be used to guide the partitioning of clock models in molecular dating (e.g., Duchêne et al. 2016; Foster and Ho 2017).

A much more complex and parameter-rich model of genomic evolution is the ‘degenerate multiple pacemaker’ model, which asserts that all loci have independent patterns of among-lineage rate variation, such that their rates are fully uncorrelated (Snir et al. 2012). This model can also be placed in the context of a Euclidean space where variables are branches in the phylogeny, such that the independence of loci means that they cannot be summarized in any lower-dimensional space (Fig. 1d). Under this condition, statistical modelling is difficult, impairing molecular dating and tests of any correlation between molecular rates and biological traits.

As an alternative to clustering into groups, loci might show relatively continuous gradients in rates. In this case, a subset of lineages might show increased variance in rates in a subset of loci. Such a scenario is perhaps the most realistic and might be apparent when rate variation is being driven by a multitude of factors. Subsets of lineages might experience co-varying rates in subsets of loci, creating “axes” of variation (Fig. 1e). Each axis can then be used to gain insight into the functions of genes that are associated with the lineages involved, and into the lineage traits that drive rate variation across the phylogeny. We call this scenario the ‘multiple local pacemaker’ model. Testing hypotheses of lineage contribution to variance along specific rate axes would offer the most complete picture of variation in rates across the genome and across a phylogeny.

We present ClockstaRX, a comprehensive tool for molecular clock visualization and hypothesis testing for genome-scale data sets. This tool is a complete rewrite with extensive improvements on its predecessor, ClockstaR (Duchêne et al. 2014), solving several difficulties associated with exploring molecular rate variation. The primary goal of ClockstaRX is to allow testing of the factors that drive rate variation, including identifying loci or branches with distinctive rates of evolution. We evaluate the performance of this tool using an extensive set of simulations and a phylogenomic data set from birds. ClockstaRX is aimed at being user-friendly and in producing data and figures that are ready for biological interpretation, further analyses, and publication. It is implemented in R and freely available with step-by-step tutorials through github.com/duchene/ClockstaRX. All of the code and results of the analyses performed in this study are available in github.com/duchene/crxTests.

## New approaches

### Collection of rates data

The basic input for ClockstaRX is an inferred species tree and a set of gene trees. The gene trees must be phylograms, with branch lengths representing expected substitutions per site. The user can choose to ignore branch lengths in the species tree. Alternatively, a species time-tree can be used as input so that the gene-tree branches are scaled into molecular rates. If a species time-tree is used, the relative branching times are assumed to be accurate.

ClockstaRX uses phylogenetic quartets to collect estimates of molecular rates. The internal branches of quartets in gene trees are used, but only where the quartet does not conflict with the species-tree topology. When a quartet in a gene tree is not present in the species tree, the information from the internal branch is not collected. In this case, the missing data for that branch can then be replaced with the mean across the branch or another imputed value that has minimal influence on the Euclidean space of the complete data set. To allow assessment of the impact of missing data across loci, ClockstaRX reports the number of times each branch in the species tree is present among the gene trees.

The quartet-based approach bypasses some potential sources of bias associated with analyses of rates, including missing taxa in some gene trees, node-density effects (Hugall and Lee 2007), and gene-tree discordance due to incomplete lineage sorting or gene-tree estimation error (Mendes and Hahn 2016). The collection of rate estimates from only the gene-tree branches that are present in the species tree places an emphasis on data reliability. However, this criterion can be overly stringent when discordance is minor, such that it comes at the expense of a loss of power and even bias from excluding the information about rates in nearly concordant tree topologies or very slowly evolving loci. Nonetheless, this form of data collection allows rate estimates to be obtained from gene trees with missing taxa, which are a common feature of phylogenomic data sets.

### Raw and weighted clocks

The whole-genome pattern of branch lengths, expected to be driven by lineage effects, can hide possible gene-specific phenomena driven by residual effects. In addition, if the input species tree is not a time-tree, the only information about gene-tree branch lengths is the phylogram, which partly depends on the time durations. To extricate the residual effects from the dominant lineage-specific signals, ClockstaRX performs two analyses by default. One is aimed at exposing lineage effects by using the raw data collected from gene trees. If a species time-tree is supplied, then the gene trees are turned to ratograms, where the original branch lengths are divided by their time durations. Another is aimed at exposing residual effects after correcting for lineage effects. This is done by transforming the data for each branch using a weight that is the relative mean length of the branch across loci (Gillespie 1989; Bedford and Hartl 2008). In the absence of a species time-tree, the transformation leads to residual phylogram branch lengths, rather than residual rates from ratograms.

### Modelling the Euclidean space of rates

ClockstaRX implements two approaches for describing the high dimensionality of rates across branches and loci. The first approach follows its predecessor, ClockstaR, in using multidimensional scaling (MDS) to map gene trees into two dimensions. This mapping depends on a matrix of pairwise distances between loci, where each pairwise distance is given by a metric that summarizes the difference between two gene trees in their branch lengths (Duchêne et al. 2014). Calculating this matrix involves a high computational cost, so ClockstaRX allows its calculation in parallel across computer cores. However, the procedure is still prohibitive for data sets with thousands of loci.

Instead of mapping gene trees using MDS, the preferred method in ClockstaRX is to model the main axes of variation in rates using principal components analysis (PCA). The *n × p* matrix used in PCA consists of the *n* loci across rows and the *p* branches of the species tree as columns. Working with this data structure is highly computationally efficient and allows formal tests of molecular clock hypotheses (see *Testing molecular clocks*).

Under this framework, each principal component can incorporate the variance in rates across all branches simultaneously. The first component represents a model of the maximal correlation across branches. The loadings of branches measure their correlation with each principal component, so it is straightforward to rank branches by their contribution to variance in each component. Similarly, the values associated with loci in each principal component indicate the relative rates of these loci on the branches with high loadings, allowing identification of loci with unusually high or low rates on those branches.

### Clustering of loci based on among-lineage rate heterogeneity

ClockstaRX identifies the number of clusters of loci, where each cluster has a distinct pattern of among-lineage rate heterogeneity. As in ClockstaR, the Partitioning Along Medoids algorithm (Kaufman and Rousseeuw 1990) is used to perform clustering, and the procedure is repeated for each of *k* number of clusters. For clustering under each value of *k*, the gap statistic of cluster dispersion is calculated. Then, the dispersion of clusters is compared with a bootstrapped sample of the data (Tibshirani et al. 2001), and the optimal number of clusters is taken as the lowest *k* that represents a peak across Gap values. An additional criterion in ClockstaRX is to select one single cluster in cases where there is a large drop in the Gap statistic from one cluster to two, which we propose as a more appropriate criterion in the scenario where clusters are poorly defined.

### Testing molecular clocks

Describing molecular clocks using PCA allows explicit tests of molecular clock hypotheses. ClockstaRX implements three types of tests, starting with a test of the degenerate multiple pacemaker model of genome evolution (Snir et al. 2012), where the null hypothesis is that of fully independent rates across loci and branches. The test statistics implemented, *φ* and *ψ*, use the magnitudes of PCA eigenvalues to estimate the overall degree of correlation between variables (Vieira 2012; Björklund 2019). The degree of covariation in the empirical data is then compared with that under permutation of samples (in this case the rate at each locus) within each variable (in this case the branches).

The test therefore assesses whether there is any covariation in rates among branches, such that dimensionality in the Euclidean space can be summarized in some lower-dimensional space. Such covariation among branches must not be confused with rate autocorrelation across branches (Kishino et al. 2001), since in ClockstaRX branches do not need to be close relatives for their rates to be correlated. A significant result from this test indicates that there is a predictable component of rate variation across lineages and loci, rejecting the degenerate multiple pacemaker model.

A second test identifies the number of pacemakers required to describe the data, and specifically the number of principal components that significantly describe variation in the data. A principal component that describes a greater proportion of the variance than the permuted sample can be considered as a variable that significantly describes evolutionary rate variation, and therefore a ‘pacemaker’. This approach is also a test of the complexity of the signal across branches and loci.

A third test identifies local pacemakers by evaluating whether each branch is significantly contributing to the signal at each principal component. This is done by testing whether the variable loadings at each principal component are greater than those under permutation. This test therefore evaluates whether specific branches have a significantly greater influence on each principal component than expected under permutation, and are therefore driving variation in evolutionary rates.

### Testing correlates of variation

ClockstaRX provides basic visualization of possible correlates of locus rates in Euclidean space. By default, the output includes the data across the first two principal components, coloured by a variety of metrics associated with the gene trees used as input. These metrics include the pattern of clustering of loci (Fig. 2; see other examples below), overall locus rate, and mean branch support per locus. Some metrics can have a strong association with evolutionary rate variation, helping to identify its causes. In addition, the user can provide any number of other variables that might be associated with the distribution of gene trees in Euclidean space, such as whether they represent coding or non-coding loci, their chromosome, or whether they include specific taxa or internal branches of interest.

**Fig. 2.**
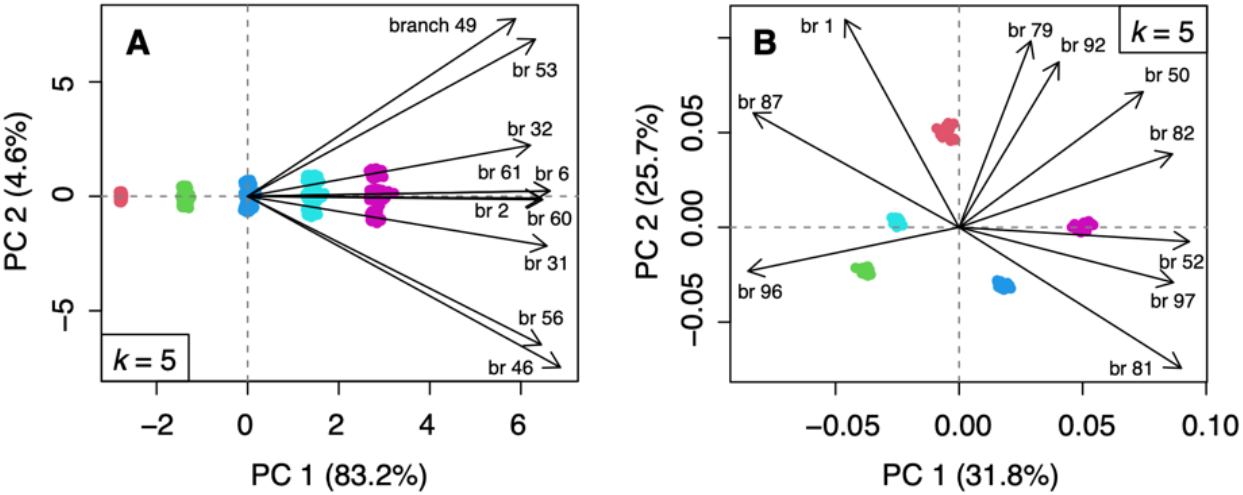
PCA plots generated by ClockstaRX from simulated phylogenomic data. Data points represent loci in a Euclidean space of rates where variables are species-tree branches, with their correlation modelled using principal components analysis. Colours indicate the clusters of loci identified. In both panels, the inferred number of clusters matched the number in the original simulation (*k* = 5). Vector arrows show the direction of increasing rates of the ten branches with the most variance, and their length is proportional to their variance across the two principal components shown. (a) Plot based on data that were simulated such that each group of loci had a distinct overall evolutionary rate, but with all groups sharing the same pattern of among-lineage rate heterogeneity (approximating the universal pacemaker model). (b) Plot based on data that were simulated such that each group of loci had a distinct pattern of among-lineage rate heterogeneity (approximating the multiple pacemaker model).

## Results and discussion

Evolutionary rate variation across taxa and genomic regions can be highly complex and therefore challenging to visualize and characterize. We present software and methods that intuitively and efficiently evaluate patterns of rate variation in data sets comprising large numbers of gene trees, allowing the identification of distinctive taxa and loci that might warrant further inquiry or special consideration in downstream analyses. The results of our simulation study, which included challenging scenarios of rate variation, allow us to provide an evaluation of our software and recommendations for its use.

### Identifying dimensional reducibility of clocks

Our extensive simulation study shows that the first test of molecular clocks described above, of the degenerate multiple pacemaker, is influenced by the simulated amounts of among-lineage rate heterogeneity and by how loci are clustered by their patterns of among-lineage rate heterogeneity (Table S1). We find that the degenerate multiple pacemaker model is nearly always rejected for data with intermediate amounts of among-lineage rate heterogeneity (96% of simulations; Fig. S2). The model was also rejected for data with negligible among-lineage rate heterogeneity when loci were simulated to be clustered by rates across all lineages (i.e., by overall rate; 92% of the time), but was rarely rejected when loci were simulated to be clustered by different patterns of among-lineage rate variation (30%).

Therefore, if genomic data contain negligible amounts of rate variation, they will appear as lacking any correlation among dimensions in the Euclidean space of rates. This means that the pattern of rate variation in non-informative data will resemble that of a degenerate multiple pacemaker (Snir et al. 2012). Dimensionality reduction was particularly difficult when among-lineage rate variation was extremely high, with the data clustered by overall rate being less dimensionally reducible than data clustered by patterns of among-lineage rate variation (26% versus 71% of simulations). Therefore, our simulations demonstrate that scenarios with extremely high or low among-lineage rate heterogeneity can be identified by ClockstaRX as lacking structure in Euclidean space.

### Identifying clusters

The number of clusters of gene trees in the Euclidean space of rates is identified most accurately by ClockstaRX when there is a single cluster, with reduced accuracy when the number of clusters is large (Fig. 3). This suggests that inferring the number of clock clusters using the technique implemented here (partitioning around medoids) is difficult compared with testing other clock hypotheses, such as the universal or degenerate multiple pacemaker models. Missing data that were introduced in simulations as caused by incomplete lineage sorting, and under conditions of high among-lineage rate heterogeneity, led to underestimates of the number of clusters.

**Fig. 3.**
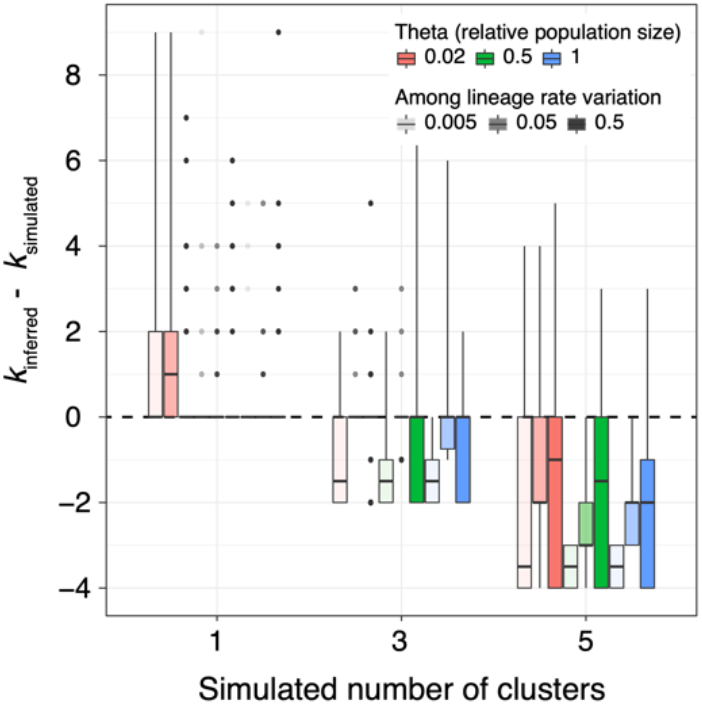
Difference between inferred and simulated number of clock clusters. The dashed line indicates the target value. The data are separated by the three simulation variables that best explain the number of clusters identified, which include the number of clusters simulated, the degree of gene-tree discordance (determined by the relative population size), and the degree of among-lineage rate heterogeneity.

Given these results, we advise users to first examine their data for highly non-clocklike loci (Vankan et al. 2022). This can be done by identifying and excluding the loci that fail Felsenstein’s likelihood-ratio test of clocklikeness, using a significance threshold that excludes only loci with severe degrees of rate variation (α = 0.001). Similarly, analyses might benefit from excluding unusually long branches, which can be done by assuming that branches follow some distribution (e.g., exponential) and excluding any that fall in the tail (e.g., *p* < 0.01; also see Mai and Mirarab 2018).

Good practice also involves verifying that loci are sufficiently variable to allow branch rates to be inferred reliably (Dornburg et al. 2019; Duchêne et al. 2022); loci that have evolved too slowly or quickly tend to yield gene trees with large estimation errors. Other factors in our simulations had relatively small impacts on the accuracy of estimates of the number of clusters (Table S1). The gap statistic implemented in ClockstaRX for the selection of the number of clusters (also proposed earlier by Duchêne et al. 2018) had greater accuracy than other existing criteria, with a slight tendency towards underestimation of *k* (Fig. S3).

### Identifying branches with high variation in rates

Branches with a strong influence on rate variation were accurately identified in our analyses of data that had been simulated so that loci shared a single pattern of among-lineage rate heterogeneity (Fig. 4). This performance was reduced when the data had multiple clusters of patterns of among-lineage rate heterogeneity, and error was further increased when large numbers of loci were included. Therefore, accurately identifying branches with high variance in rates is likely to be more difficult for data sets with pronounced clustering of loci. These might be loci that have intrinsic differences in their evolutionary history, such as coding vs noncoding DNA or loci on different chromosomes, and suggest that a subset of taxa follow a process comparable to a universal pacemaker model. In every instance, the presence of large numbers of branches that were highly influential in driving patterns of rate variation led to increased accuracy in the proportion of such branches being identified.

**Fig. 4.**
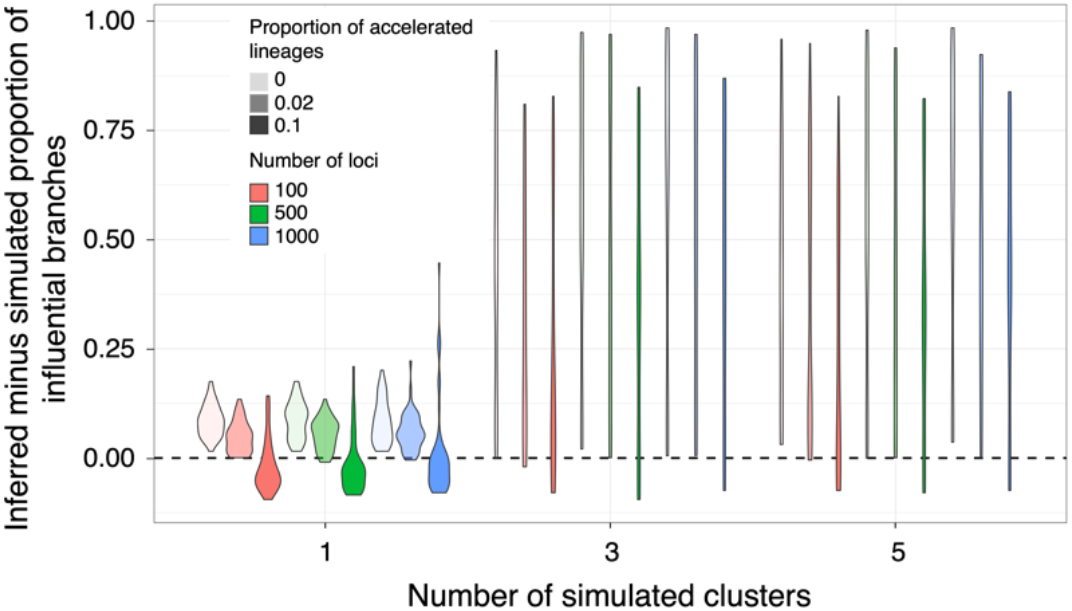
Difference between inferred and simulated proportions of influential branches. The grouping of data corresponds to the most influential variables, including the difference between inferred and simulated proportion of influential branches, was better explained by the simulated proportion of branches with rate acceleration.

### Case study: Avian genomes

An analysis of a subsample of 200 introns and exons sampled from 42 species across nearly all avian orders (Jarvis et al. 2014) showed that the loci formed two distinct clusters on the basis of their patterns of rate variation (Fig. 5a). The two clusters corresponded almost exactly to introns and exons (Fig. 5e), suggesting that these two data types have distinct patterns of among-lineage rate heterogeneity. Consistent with the results of our simulation study, the clusters were identified with most confidence after branches were weighted to remove lineage effects (Fig. S4).

**Fig. 5.**
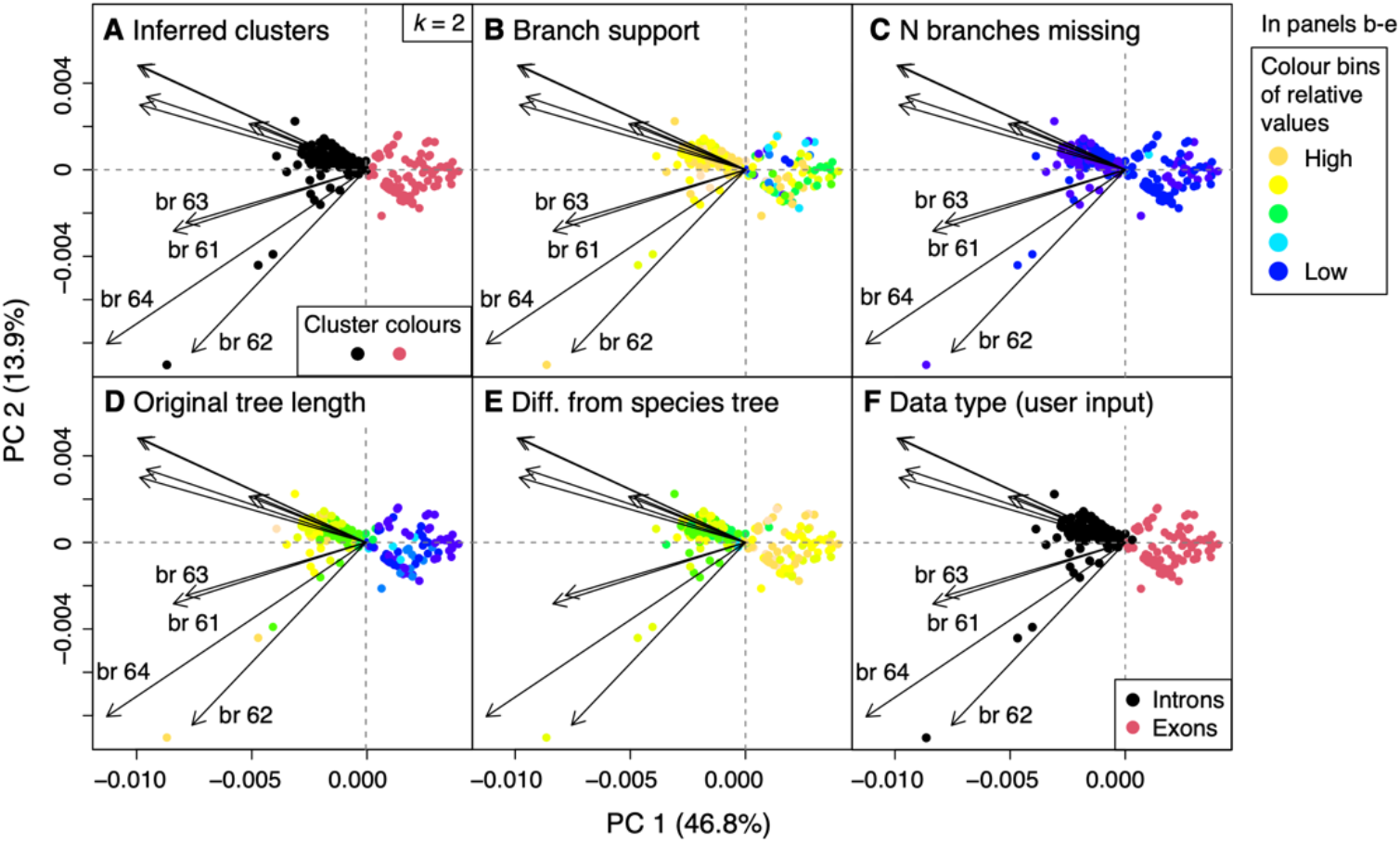
ClockstaRX results from an analysis of 200 loci sampled across avian orders. Data points represent loci and are coloured in panels (a) to (e) by variables that are by default inferred or extracted by ClockstaRX. In panel (f), loci are coloured by user-input data, which in this case is data type. In panels (b) to (e), warmer colours indicate greater relative values, such that exon loci are seen to have lower branch support (b), greater numbers of missing branches (c), lower tree lengths (d), and greater unweighted Robinson-Foulds topological distances from the species tree (e). While the top ten branch-variable vectors are shown, only those leading to passerine lineages are labelled (br 61 to br 64).

Tests of individual principal components provided additional information about the drivers of heterogeneity in the data. A single principal component explained around half of the variance in rates across loci (46.8%), suggesting that a simple process, such as that driving relative locus rates, is the dominant driver of heterogeneity. In support of this, tests only supported two principal components as explaining a significant amount of variation in the data (*P* < 0.001; Fig. S4). This can also be seen in the comparable directions of branch-variable vectors (Fig. 5), which indicate an overall tendency of higher substitution rates in introns.

The separation between introns and exons reflects the well-known difference in selective constraints between the two data types (e.g., Hughes and Yeager 1997; Laroche et al. 1997). The dominant distinction between introns and exons appears at first to be relative substitution rates across loci, following a universal pacemaker. The lack of an overlap in rates between introns and exons seems surprising; the varying degrees of selection in introns can be expected to somewhat overlap the amount of selection found of exons. Null expectations of various degrees of selection across clusters or data types could be tested using simulations and used to assess the crucial role of introns in gene regulation (Chorev and Carmel 2012; Rose 2019). Specifically, the high evolutionary rates of introns might be driven by the fast-evolving nature of adaptive gene expression (Ghalambor et al. 2015; Sands et al. 2021), rather than simply reflecting the underlying rates of neutral mutation.

The support for a second principal component in these data (explaining 13.9% of the variance) suggests that other processes have driven variation in evolutionary rates. These additional processes were likely to be driven by the traits found in specific lineages, as observed in the larger vectors observed for some branches (Fig. 5). Test of individual branches for significant contribution to rate heterogeneity in each of the two principal components confirmed that the rate heterogeneity was due to a subset of branches. Nonetheless, as found in simulations, given the signal of clustering in the data into introns and exons, there might be high stochastic error in the identification of branches as influential.

Passerine lineages had the greatest variance in rates along both principal components (branch vectors labelled br 61 to br 64 in the figures). This is consistent with the hypothesis that passerines have undergone distinct evolutionary trajectories in their genomes, brain development, locomotion, and life history (Zhang et al. 2014). Additional analyses of larger portions of genes and individual data types could be used to identify the gene families that have driven the explosive diversification of this group of birds. Candidate genes include those associated with hormone signalling (Schlinger et al. 2022), growth hormone family (Yuri et al. 2008; Feng et al. 2020), cognition (Gesicki 2022), and immune system (O’Connor et al. 2016). Non-coding regions might also reveal an overall higher rate of mutation in passerines compared with other avian taxa, which could be associated with their shorter generations (Cooney et al. 2020).

### Limitations

ClockstaRX excludes any quartets in the gene trees that are not present in the species tree, with the aim of improving the accuracy of rate estimates. This comes at the cost of losing some data and a possible loss of power, and even bias towards longer branches that can be estimated reliably. Nonetheless, length estimates of short branches and from loci with low rates are often associated with low accuracy and high uncertainty, such that their exclusion is unlikely to be a major source of bias.

Some signal might also be lost when there is a large amount of phylogenetic estimation error. Before analysis in ClockstaRX, the amount of error can be reduced by taking a more aggressive approach to data filtering. For instance, whole loci can be excluded if their branch supports are too low (e.g., mean < 0.9) or if the substitution model is deemed inadequate by software such as PhyloMAd (D.A. Duchêne et al. 2018) or IQTREE2 (Bui et al. 2020); individual taxa can be excluded if they are associated with large amounts of topological uncertainty, using software such as RogueNaRok (Aberer et al. 2013) or TreeShrink (Mai and Mirarab 2018); alignment segments that are likely to be misaligned can be identified and removed using software such as PREQUAL (Whelan et al. 2018).

Ideally, the source of the problem should be found, and can range from the rate of evolution of the region of interest, sequencing quality of samples, alignment, or the model used for phylogenetic inference. In particular, methodological artifacts associated with alignment are worth studying further (Warnow 2021), such as varying amounts sequencing quality, sequence conservation, and selection (e.g., Vakhrusheva et al. 2013; Frith et al. 2021). Since there is a large range of possible sources of misleading data, it is perhaps most appropriate to use automated methods of data filtering in phylogenomic data, combined with some visual inspection of the data.

Furthermore, discordance between the gene trees and the species tree will occur under conditions of high incomplete lineage sorting. ClockstaRX is therefore particularly suited for data sets where loci share a similar or identical topology, such as when population sizes have been small relative to the waiting times between divergence events. When these conditions are strongly violated, it is perhaps most appropriate to focus on genes of interest or exclusively on lineage effects. Nonetheless, the partial filtering in ClockstaRX maximizes the amount of reliable data, and missing data do not appear to affect the inferred location of loci in space (Figure 5).

Despite the overall good performance of ClockstaRX in analyses of simulated data, some signal is likely to be lost in visualization and analysis of dimensionally reduced data. This is also a common difficulty when summarizing the variation in phylogenetic signals across loci (D. A. Duchêne et al. 2018; Smith 2021). Data loss and distortion might have contributed to the difficulties that we observed for data that had been simulated with extremely low or high degrees of among-lineage rate heterogeneity. Similarly, factors other than those explored in our simulations could be influential, such as the diversification process, model of rate variation across branches (e.g., autocorrelated or following a parametric distribution), or distribution of loci across clusters. Therefore, analyses of empirical data will benefit from careful data curation and exploration before any inferred patterns in evolutionary rates are interpreted as having a biological basis.

### Conclusions

Evolutionary rates are fundamental biological quantities and can be understood through detailed analysis of phylogenomic data. ClockstaRX provides a user-friendly tool for achieving comprehensive description of molecular rates in phylogenomic data, shedding light on the drivers of evolutionary rates and aiding the construction of models for molecular dating. The formal tests implemented in ClockstaRX allow users to identify the lineages and loci that significantly co-vary in their evolutionary rates. The default visualizations produced by the software facilitate the assessment of any overarching patterns in rates. Therefore, a natural application of ClockstaRX is the incorporation of biological trait and gene function data, aimed at explaining the molecular clock phenomena that are found. By placing molecular clocks into a comparative analytical framework, ClockstaRX aims to expand our understanding of evolutionary rates and timescale across genes and lineages in the Tree of Life.

## Materials and methods

We generated synthetic data under a broad range of simulation scenarios to allow us to examine the performance of ClockstaRX. Specifically, we examined whether factors that often vary across empirical data sets can affect the detection of any lower dimensionality in the Euclidean space of rates, and the detection of clusters of loci and branches that have a strong influence on patterns of rate variation. In each simulation, we generated a species tree under a birth-death process (*N* = 50 tips; age = 50 time units; *λ* = 0.5; *μ* = 0.1) using *TreeSim* (Stadler 2011). Gene trees were then simulated as embedded in each species tree under the multispecies coalescent with a constant population size as implemented in *phybase* (Liu and Yu 2010). Gene-tree discordance was assumed to arise exclusively from incomplete lineage sorting (*θ* = 0.02, 0.5, or 1).

Rates of molecular evolution were simulated for each gene tree under a white-noise molecular clock model (Lepage et al. 2007), as implemented in *NELSI* (Ho et al. 2015). Clock models varied across scenarios in their mean substitution rate (mean = 0.01, 0.05, or 0.1 substitutions per site per time unit) and the extent of among-lineage rate heterogeneity (mean standard deviation = 0.005, 0.05, and 0.5).

Other parameters that we varied across our simulations were the number loci sampled (100, 500, or 1000) and the number of clock clusters (*k* = 1, 3, or 5). We allowed two possible types of clock clustering of loci: by overall evolutionary rate (affecting all branches), or by an individual pattern of among-lineage rate heterogeneity in each cluster. In the first, clusters were numbered 1 to *k* and overall rates in each cluster were multiplied by the cluster identity (e.g., loci in cluster 5 had rates five times those in cluster 1).

The second method of clustering simulated a single set of branch rates for all loci within each cluster. In this scenario, each branch rate in each locus also had normally distributed variation under a standard deviation of 0.1 times the original branch rate. A last form of variation involved highly influential branches with accelerated rates. In each scenario, a proportion of branches (0, 0.02, 0.1) were given rates that were five times their original values. Given that the proportion and location of branches could vary widely, our analyses focused on the accuracy in identifying a given portion of accelerated lineages, rather than the specific identities of such lineages.

The gene trees produced by each simulation were processed in ClockstaRX to extract three types of information: dimensional reducibility as tested by the *φ* and *ψ* tests, the number of clusters *k* in the data, and the proportion of branches with accelerated rates due to dominant lineage effects. We tested the simulation variables individually in linear regression for their ability to explain the accuracy of ClockstaRX inferences.

To demonstrate the usage of ClockstaRX in empirical data, we analysed a sample of introns and exons from 42 bird species, representing most extant orders (Jarvis et al. 2014; Jarvis et al. 2015). We first estimated the maximum-likelihood tree for each alignment under the selected GTR+I+F+Γ+R model as implemented in IQ-TREE2 (Bui et al. 2020). We then randomly sampled 100 introns and 100 exons, and used the dated species tree from the original study for analysis in ClockstaRX.

## Acknowledgements

This work was supported by funding from a European Research Council Marie Sklodowska-Curie fellowship to D.A.D. (H2020-MSCA-IF-2019-883832), and the Australian National Health and Medical Research Council awarded to S.D. (APP1157586).

