## Supplementary material for "ClockstaRX: testing molecular clock hypotheses with genomic data": Table S1

**Table S1.** P-values of regression analyses testing the association between each of the simulation variables (columns) and eight estimates from ClockstaRX (rows).

| | Theta (relative population size) | Number of loci | Among-lineage rate variation | Proportion of lineages with accelerated rate | Number of clock clusters | Mean overall rate | Clustering type (overall rate or clock) |
|---|---|---|---|---|---|---|---|
| $k$ | <0.0001 | 0.0312 | 0.0105 | 0.1153 | 0.1718 | 0.1131 | 0.0027 |
| $k_{inferred}$ - $k_{simulated}$ | <0.0001 | 0.0946 | 0.0472 | 0.2217 | <0.0001 | 0.2189 | 0.0201 |
| $k$ (weighted clock data) | <0.0001 | 0.0024 | 0.0006 | 0.0420 | <0.0001 | 0.9807 | 0.1987 |
| $k_{inferred}$ - $k_{simulated}$ (weighted clock data) | <0.0001 | 0.0127 | 0.0048 | 0.0955 | <0.0001 | 0.9842 | 0.2918 |
| Inferred proportion of influential lineages | 0.0435 | <0.0001 | 0.0028 | 0.0651 | <0.0001 | 0.7205 | 0.0073 |
| Inferred minus simulated proportion of influential lineages | 0.0469 | <0.0001 | 0.0033 | <0.0001 | <0.0001 | 0.7246 | 0.0082 |
| $\phi$ statistic | 0.1717 | 0.0691 | <0.0001 | 0.0845 | 0.0658 | 0.5646 | 0.0191 |
| $\psi$ statistic | 0.1798 | 0.0640 | <0.0001 | 0.0830 | 0.0669 | 0.5632 | 0.0200 |

**Figure S1.** Workflow for ClockstaRX analyses. Main function names are shown below steps in italics.

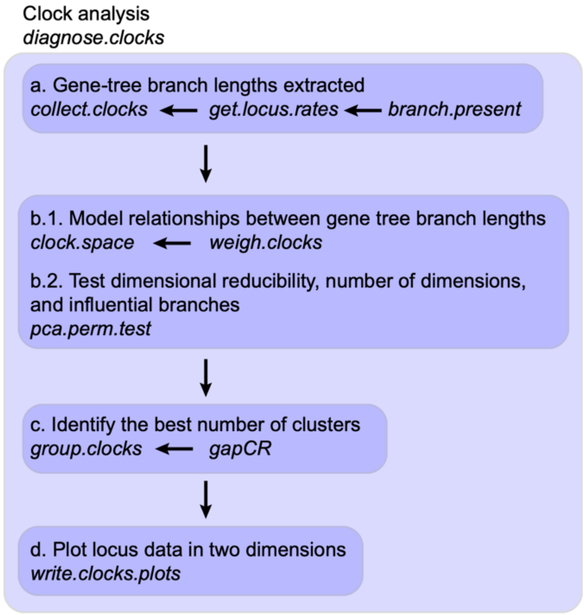

**Figure S2.** Inferences from ClockstaRX from data simulated under a range of scenarios of evolutionary rates across loci and lineages. Inferences of interest include (a) number of clusters inferred, and (b) inferred proportion of influential branches in the data.

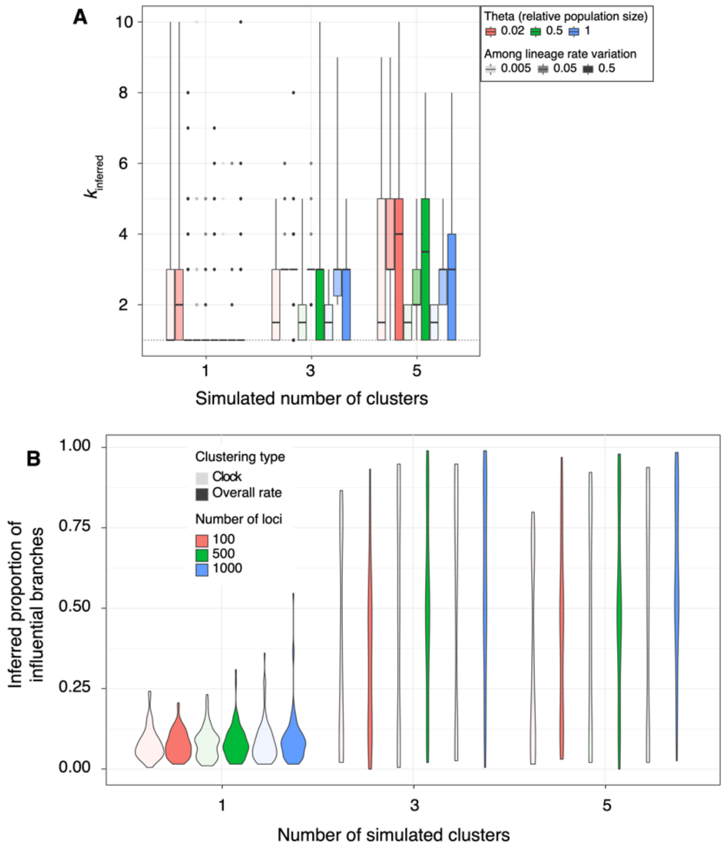

**Figure S3.** Results of the *Φ*-test of dimensional reducibility across simulations, focusing on the two simulation factors that best explained the test results. Intermediate amounts of among-lineage rate variation show the best performance, frequently rejecting the null hypothesis of a lack of correlation in the data (*P*-value < 0.01). Extremely high or low values of among lineage rate variation show generally poor performance. One exception is that of scenarios of clustering by patterns of among-lineage rate variation and low overall variance in rates, which leads to frequent rejection of the null.

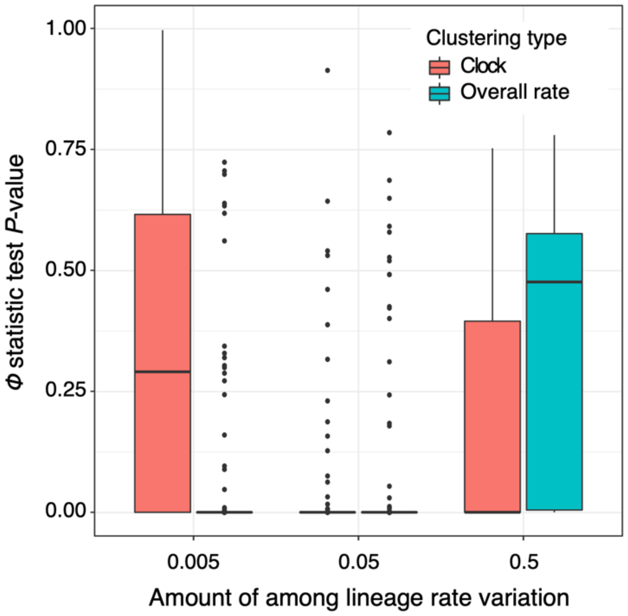

**Figure S4.** Comparison of metrics for selecting *k*, showing the number of simulations (y-axis) in which each difference between inferred and simulated *k* was found (x-axis). The error scores shown in the legend are the sum of the absolute x-axis values for each metric, such that increasing values are equivalent to a greater departure on average from the simulated *k*.

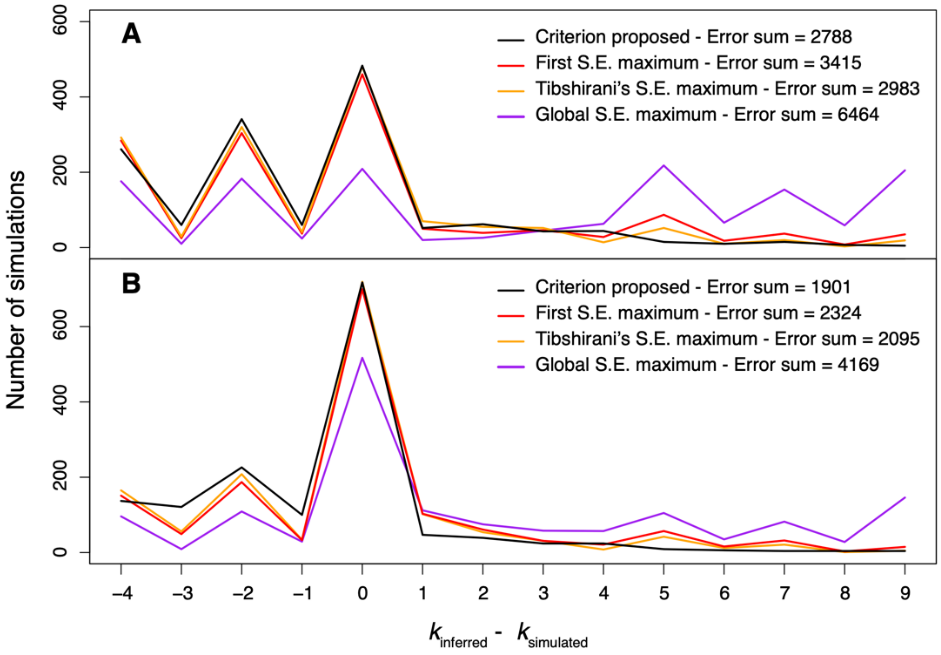

**Figure S5.** Further details of the ClockstaRX output, showing results for the example data set of introns and exons across avian orders.

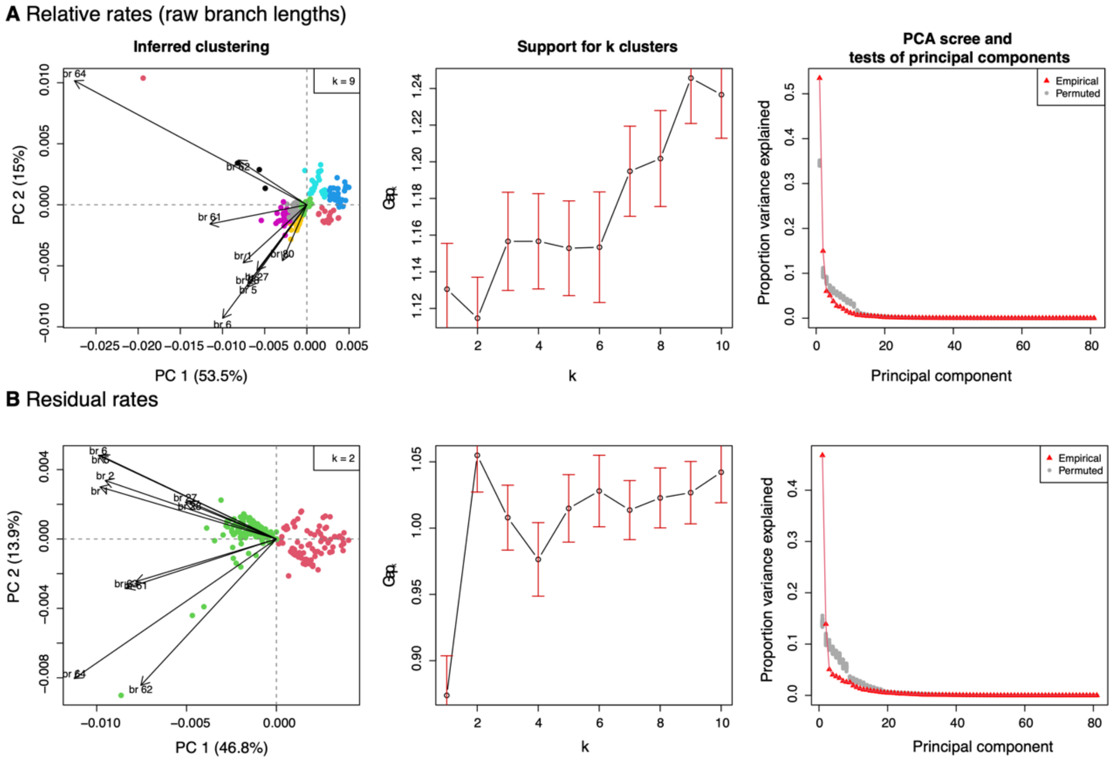
